# Controller for microfluidic large-scale integration

**DOI:** 10.1101/188615

**Authors:** Jonathan A White, Aaron M Streets

**Affiliations:** Department of Bioengineering University of California, Berkeley, Berkeley, CA 94720; Chan Zuckerberg Biohub, San Francisco, CA 94158

**Keywords:** python, Arduino shield, open source hardware, solenoid valve, microfluidics, multilayer soft lithography

## Abstract

Microfluidic devices with integrated valves provide precise, programmable fluid handling platforms for high-throughput biological or chemical assays. However, setting up the infrastructure to control such platforms often requires specific engineering expertise or expensive commercial solutions. To address these obstacles, we present a Kit for Arduino-based Transistor Array Actuation (KATARA), an open-source and low-cost Arduino-based controller that can drive 70 solenoid valves to pneumatically actuate integrated microfluidic valves. We include a python package with a GUI to control the KATARA from a personal computer. No programming experience is required.

---

**Specifications Table.**
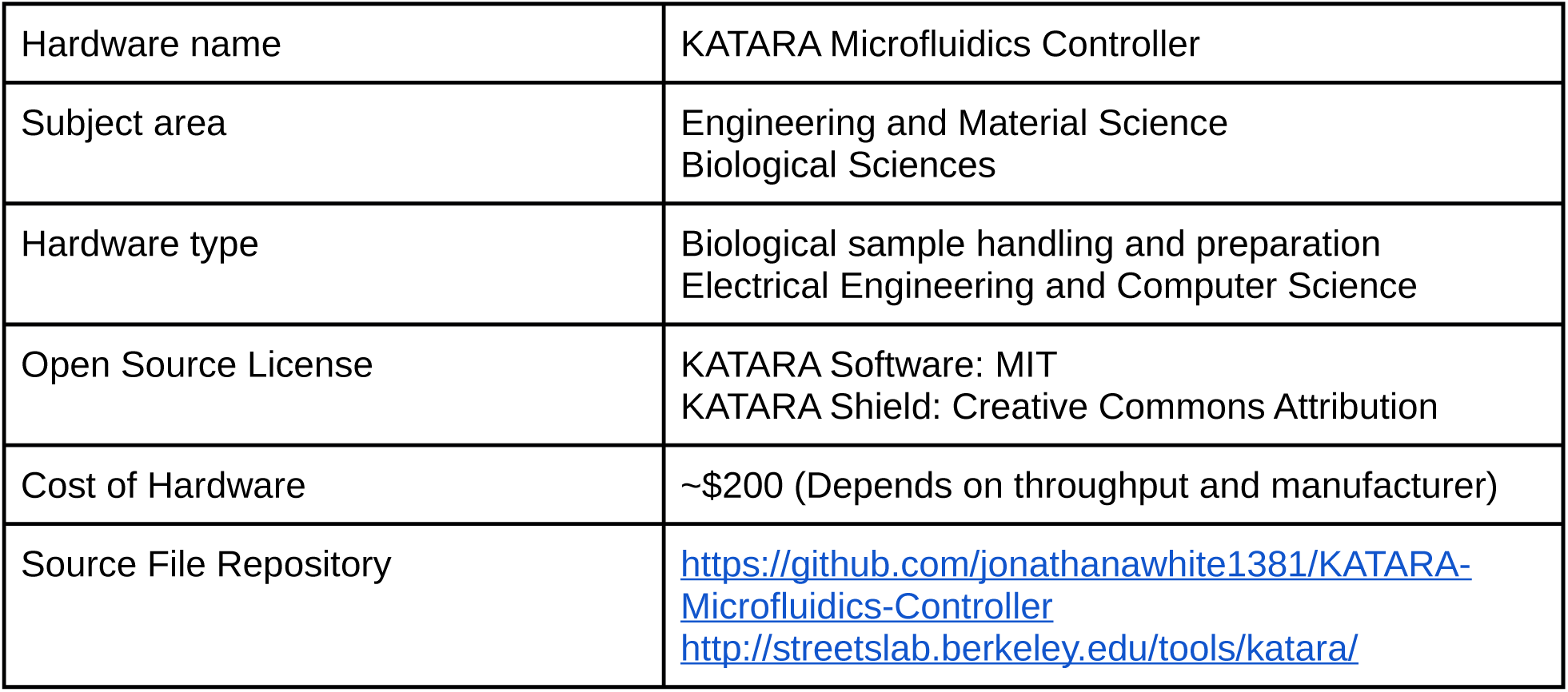

|  |  |
| --- | --- |
| Hardware name | KATARA Microfluidics Controller |
| Subject area | Engineering and Material Science<br>Biological Sciences |
| Hardware type | Biological sample handling and preparation<br>Electrical Engineering and Computer Science |
| Open Source License | KATARA Software: MIT<br>KATARA Shield: Creative Commons Attribution |
| Cost of Hardware | ~\$200 (Depends on throughput and manufacturer) |
| Source File Repository | https://github.com/jonathanawhite1381/KATARA-Microfluidics-Controller<br>http://streetslab.berkeley.edu/tools/katara/ |

## 1. Hardware in Context

Microfluidic Large Scale Integration uses micromechanical valves integrated into silicone fluidic circuits to precisely and discretely manipulate picoliters to nanoliters of liquid. This allows scientists to perform quantitative biology experiments on microfluidic devices in parallel, analogously to how integrated circuits use transistors to perform electronic computations in parallel [1-3]. Microfluidic devices with integrated valves have been developed for many applications including protein crystallization screens [4], single molecule conformation experiments [5], transcription factor binding affinity assays [6], cell culture assays [7, 8], digital PCR [9], sandwich immunoassays [10, 11], and single cell genomics [12, 13]. A commonly used micromechanical valve, known as the Quake valve, uses pneumatically actuated control channels to pinch off adjacent flow channels by deforming the interstitial wall [1] (Figure 1). This type of microfluidic valve can be manufactured in dense arrays using multilayer soft lithography [2, 14]. A major barrier to entry for using microfluidic devices with integrated valves can be implementing control software and electronics. We aim to lower this barrier by introducing a Kit for Arduino-based Transistor Array Actuation (KATARA). The KATARA microfluidics control system is an open-source and low-cost platform for writing and running procedures that actuate solenoid valves.

**Figure 1.**
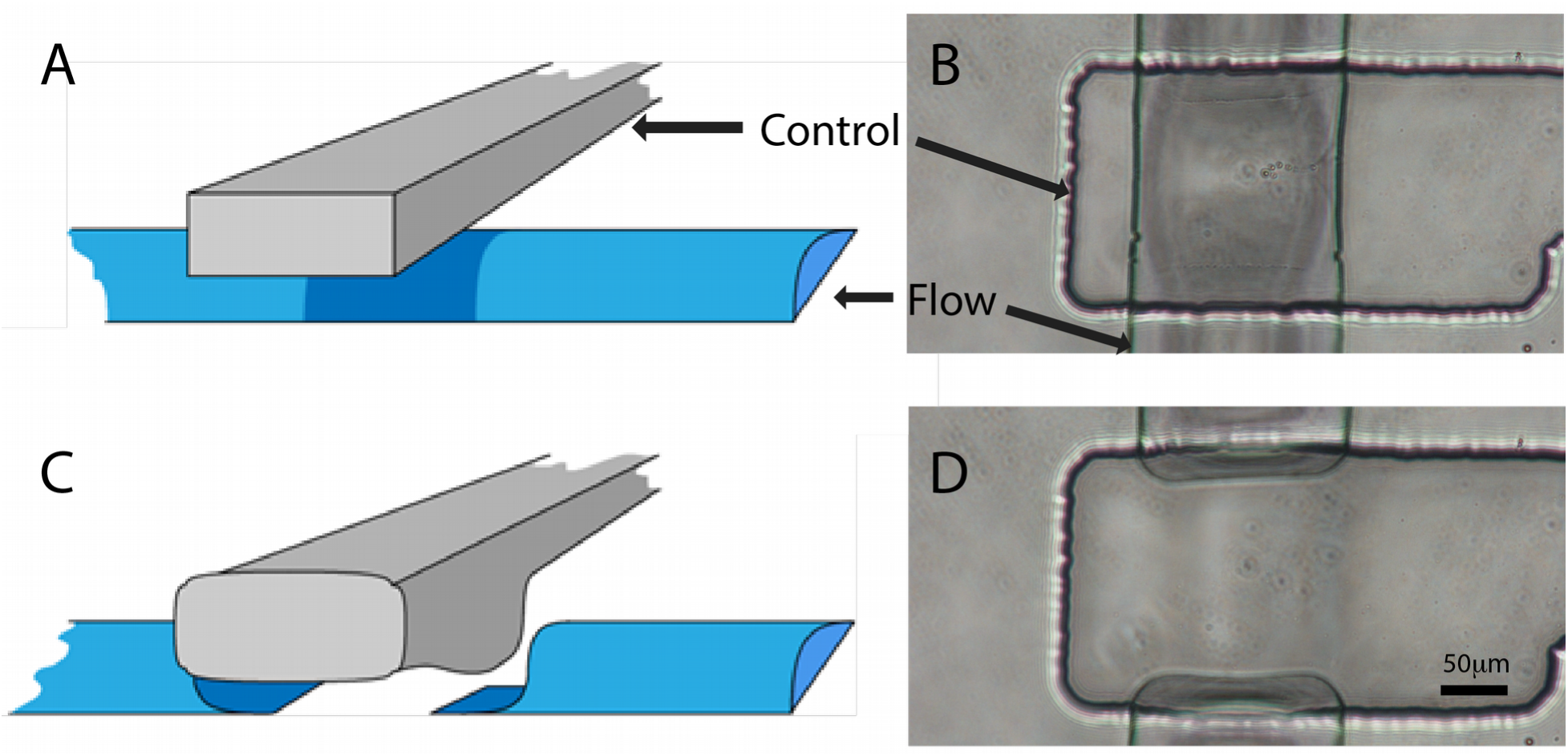
Cartoons and photographs of an integrated microfluidic valve in the open (A, B) and closed (C, D) position.

## 2. Hardware Description

The KATARA microfluidics control system includes a shield for the Arduino Mega microcontroller that drives up to 70 solenoid valves (Figure 2), Arduino firmware, and a python package with a graphical user interface (GUI). The package provides an interface to control Arduino-KATARA shield assemblies with python programs. The GUI allows users to actuate individual valves and three-valve peristaltic pumps [1] either manually or in any arbitrary automated sequence without any programming: users can create and edit protocols for pumping and actuating valves while iterating over loops (Figure 3). These protocols can be saved and loaded as custom buttons to build controls for any microfluidic device (Figure 3). The KATARA python software and firmware use Arduino digital logic pins 0 and 1 to communicate, leaving 68 pins available to control valves. To use all 70 control lines on the shield, the Arduino must be programmed directly. Solenoid valve lead wires attach to (+) and (−) terminal pairs on the shield labeled 0-69 as they are referenced in the Arduino firmware and python software. Note that lines 54-69 are labeled A0-A15 on the Arduino (Figure 2).

**Figure 2.**
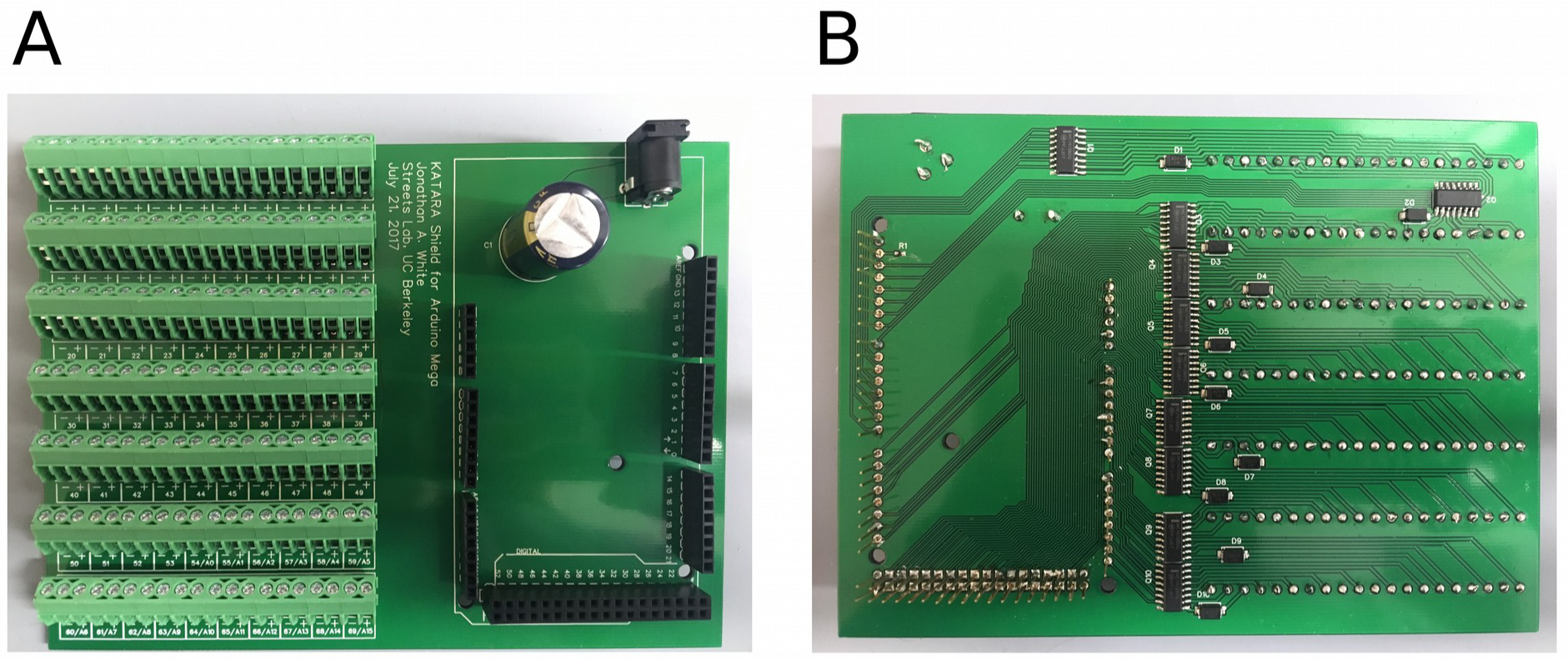
Photograph of the top (A) and bottom (B) side of an assembled KATARA shield. The top side of the KATARA shield has stackable headers, a power supply jack, and terminal blocks to attach solenoid valves. The bottom side has amplifying circuitry and the stackable header pins that plug into an Arduino Mega.

**Figure 3.**
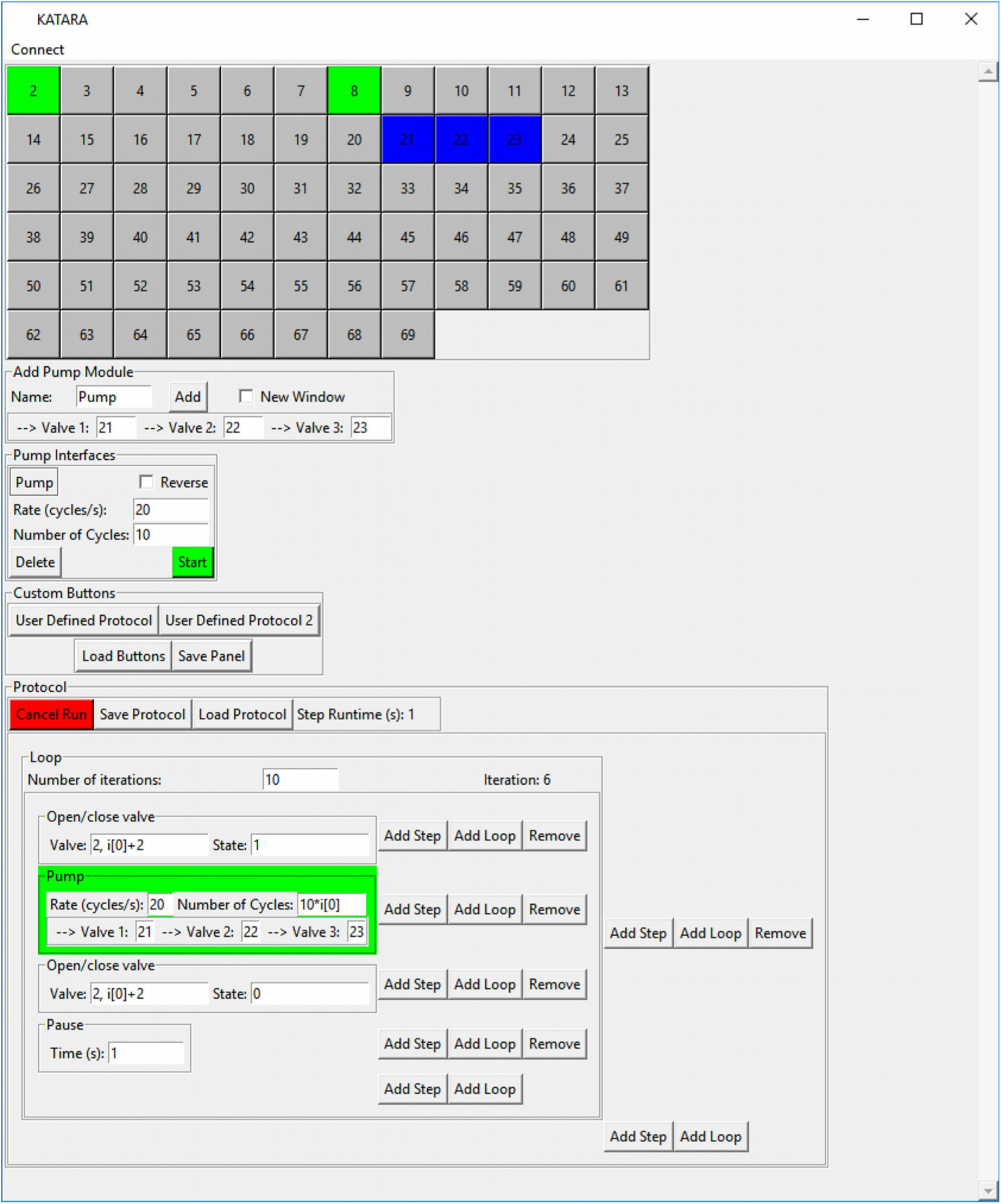
A screenshot of the KATARA GUI which has interfaces to actuate each individual valve; control peristaltic pumps; create, edit, run and save protocols; and load saved protocols as user defined buttons.

The KATARA shield is based on the open-source USB microfluidics controller designed by Rafael Gómez-Sjöberg [15], a circuit board that amplifies digital signals from a USB IO card to control 24 valves. By choosing surface-mount components, replacing the IO card with an Arduino, and streamlining valve connections, the KATARA shield can control nearly three times as many valves at a lower cost. Other available microfluidic device controllers include the Wago controller (which is thoroughly described by Brower et al. [16]), the URMC32 digital relay from numato, the Strey Lab shield for Arduino Uno that controls valves and pressure sources for eight pneumatic lines [17], the Li lab smartphone controller [18], the Maerkl lab controller [19], and a commercial device from Elveflow that can control up to sixteen valves. The KATARA software can be extended to interface with these and other microfluidic controllers (see supplement).

Researchers who use the KATARA microfluidics control system will do so because it offers:

- A low-cost circuit to control up to 70 Solenoid valves,
- A python package that allows users to control the circuit board with python programs,
- A comprehensive GUI to write and share automated protocols for experiments: no programming experience is required.

## 3. Design Files

**Table 1.** Design Files.

| File | Description |
| --- | --- |
| KATARA_Shield_EASYEDA_Footprint.json | A source file for the KATARA shield footprint editable in EASYEDA. |
| KATARA_Shield_EASYEDA_schematic_source.json | A source file for the KATARA shield schematic editable in EASYEDA. |
| KATARA_Shield_Gerbers.zip | The gerber files for manufacturing the KATARA shield. |
| Centroid_File.csv | Gives the locations of each component on the KATARA shield for assembly by pick and place machine. |
| main.py | Main script to run the KATARA GUI. |
| USB_GUI.py | Base class for graphical user interfaces that connect to USB devices. |
| KATARA_GUI.py | Contains classes for running the KATARA GUI and pump interfaces. |
| Protocol_Tools.py | Contains classes for implementing protocol interfaces in the GUI. |
| Step.py | Base class for protocol steps. |
| StepDerivatives.py | Derived step classes for pausing, pumping and actuating valves. |
| no_wait_Dialog.py | Contains modified version of the Tkinter dialog window class that does not pause the running thread. |
| LabelEntry.py | A class for drawing and referencing labeled text entry bars. |
| KATARA.ico | The KATARA icon. |
| ValveController.py | Contains ValveController and peristalticPump classes (described in supplement section 4). |
| KATARAValveController.py | Contains KATARAValveController and KATARAPump classes for sending serial signals to Arduinos running the KATARA firmware: for use by programmers and used internally in the GUI. |
| Config.py | Contains variables that GUI related classes inside different files share. |
| KATARA_Firmware.ino | The program that allows Arduinos to interpret and execute serial commands set from the KATARAValveController and KATARAPump classes. |

## 4. Bill of Materials

**Table 2.** Bill of Materials

| Designator | Component | Number | Cost Per Unit | Total Cost | Supplier | Manufacturer Part Number | Material type |
| --- | --- | --- | --- | --- | --- | --- | --- |
| Q1-Q10 | Bipolar (BJT) Transistor Array 7 NPN Darlington 50V 500mA Surface Mount 16-SOIC | 10 | \$0.45 | \$4.51 | Digikey | MC1413BDG | Semiconductor |
| D1-D10 | Zener Diode 22V 1W ±5% Surface Mount SMA | 10 | \$0.38 | \$3.84 | Digikey | SMAZ22-13-F | Semiconductor |
| P1 | Conn PWR Jack 2.5X5.5MM Solder | 1 | \$0.64 | \$0.64 | Digikey | PJ-102B | Metal/Polymers |
| MP1-MP7 | CONN TERM BLOCK 45DEG 10PS 3.5MM | 14 | \$1.99 | \$27.86 | Digikey | 1989036 | Metal/Polymers |
| R1 | RES SMD 10K OHM 5% 1/16W 0402 | 1 | \$0.10 | \$0.10 | Digikey | RC0402JR-0710KL | Composite |
| C1 | CAP ALUM 1000UF 20% 35V RADIAL | 1 | \$1.15 | \$1.15 | Digikey | EEU-FC1V102S | Metal |
| U29 | Arduino MEGA Stackable Header Kit | 1 | \$1.50 | \$1.50 | Itead | IM120531023 | Metal/Polymers |
| | AC/DC DESKTOP ADAPTER 24V 120W | 1 | \$96.88 | \$96.88 | Digikey | PSA120U-240L6 | Metal/Polymers/Semiconductor |
| | Arduino Mega 2560 Rev3 | 1 | \$45.95 | \$45.95 | Arduino | A000067 | Semiconductor |
| | Printed Circuit Boards* | 5 | \$4.98 | \$24.90 | EasyEDA | N/A | Metal/Polymers |
| | Solder Stencil* | 1 | \$13 | \$13 | EasyEDA | N/A | Metal |
| | Solder Paste* | 1 | \$15.95 | \$15.95 | Digikey | SMD291SNL | Metal |

\* These items are necessary for do-it-yourself assembly, but they will be included as part of commercial assembly services.
† Components with reference designators should be included in the bill of materials submitted to PCB assemblers.

## 5. Build Instructions

### 5.1 Hardware Design

The KATARA shield extends the Arduino Mega by amplifying signals from each of its digital pins to drive a solenoid valve. To do this, it uses ten array packages each containing seven Darlington pair amplifier circuits (Figure 4). Darlington amplifiers offer no performance advantage over simple transistor amplifiers for our application, but Darlington array packages reduce the cost and number of components necessary to build the board. The Darlington arrays include a shared flyback pin that is connected to the source voltage across the cathode of a Zener diode with 22 V breakdown voltage (Figure 4): placing Zener diodes here forces solenoid valves to close faster and ensures that the voltage across the amplifier will never exceed its 50 V rating when driving 24 V valves. The KATARA shield can also drive valves operating at voltages less than 24V; if choosing valves other than the Pneumadyne S10MM-31-24-2 to use with the KATARA shield, choose a power supply at the proper voltage that is rated to put out enough current to drive all attached valves and ensure that the average power dissipated through a valve does not exceed its maximum operating power. To ensure the power dissipated across the Zener diodes stays within specifications, limit the number of times any individual Darlington array package closes valves per second to

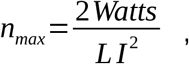

where *L* is the inductance of the valve and *I* is the current it conducts while energized. For the Pneumadyne S10MM-31-24-2, *n_max_* is greater than 100,000, but exceeding *n_max_* may be a concern with solenoid valves that draw more power. The KATARA shield also includes a 10 kΩ resistor, which connects the Arduino and external supply grounds to establish a reference for the control circuit while shielding the Arduino and the computer from current spikes, and a 1 mF capacitor, which ensures enough energy is always on hand to open solenoid valves quickly.

**Figure 4.**
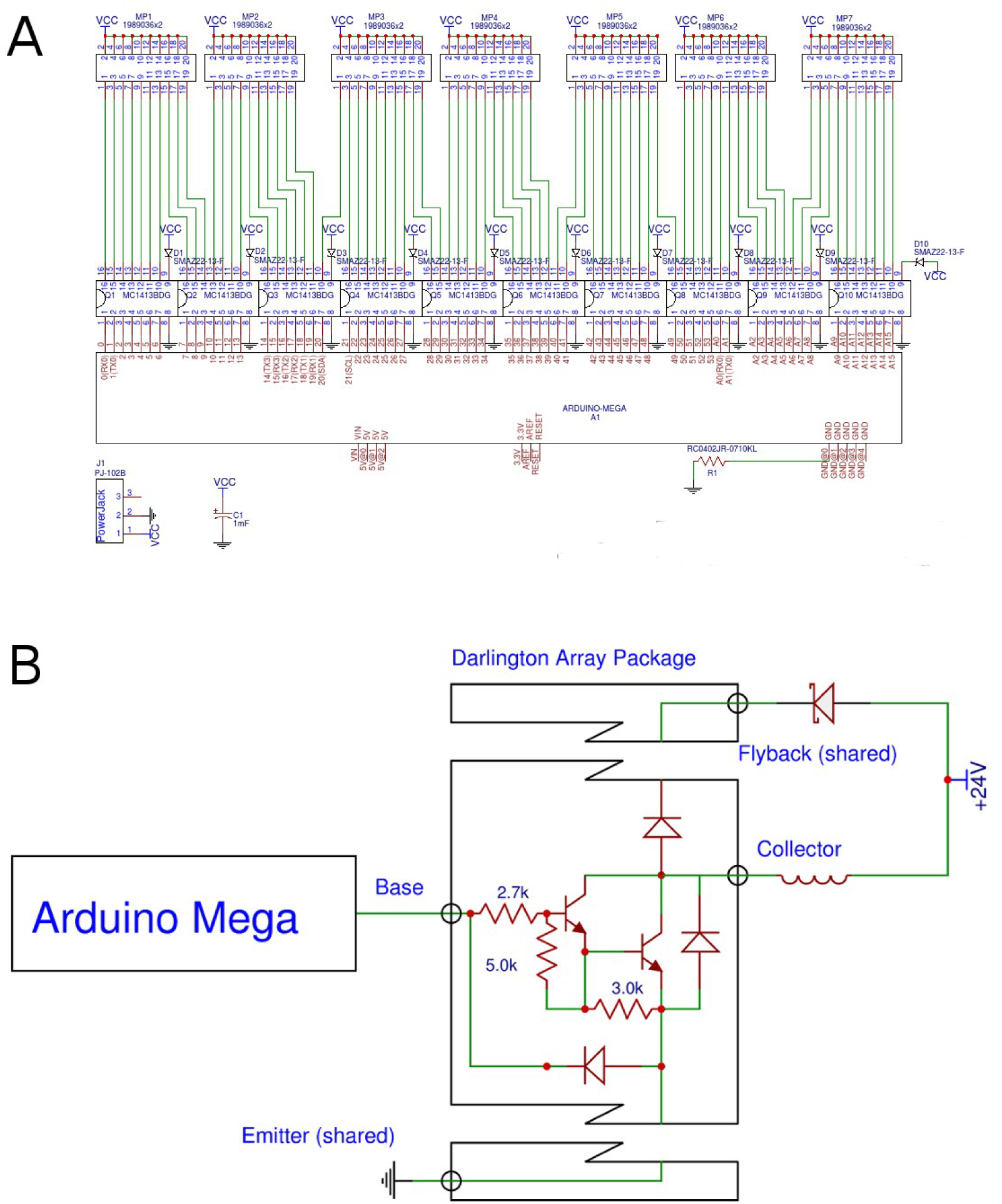
(A) Schematic of the KATARA shield. (B) Schematic of a single channel of KATARA shield.

### 5.2 Hardware Assembly

One may order assembled KATARA shields from PCB manufacturers by submitting the gerber files, centroid files, and bill of materials. To assemble the KATARA shield yourself, order an unassembled board and use the following procedure:

1. Apply solder paste to the surface mount pads on the bottom side of the board with the solder stencil.

- Reference [20] is a good tutorial on how to apply solder paste with solder stencils.
- If solder paste is accidentally applied between contact pads, surface tension will remove the connections during reflow when it pulls molten solder onto each pad.
2. Use tweezers to place the surface mount resistor (R1), transistor arrays (Q1-Q10), and Zener diodes (D1-D10) with the correct orientation.

- The white bar on the Darlington array should be oriented with the white circle in the corner of the array outline on the PCB.
- The white line printed on the diode should be oriented with the white line in the outline of the diode on the PCB.
3. Reflow solder:

- This can either be done with a reflow oven using the recommended heating profile [21] or by heating the board on a hot plate from room temperature until the solder melts at 220°C [22] to approximate the recommended heating profile and limit heat shock to the components. Be sure to perform this step in a well ventilated area.
4. Remove any excess solder that might short adjacent pads with solder wick and a soldering iron.
5. Join seven pairs of ten-position Phoenix Contact block terminals with interlocking sides to produce seven twenty-position block terminals (Figure 5).

**Figure 5.**
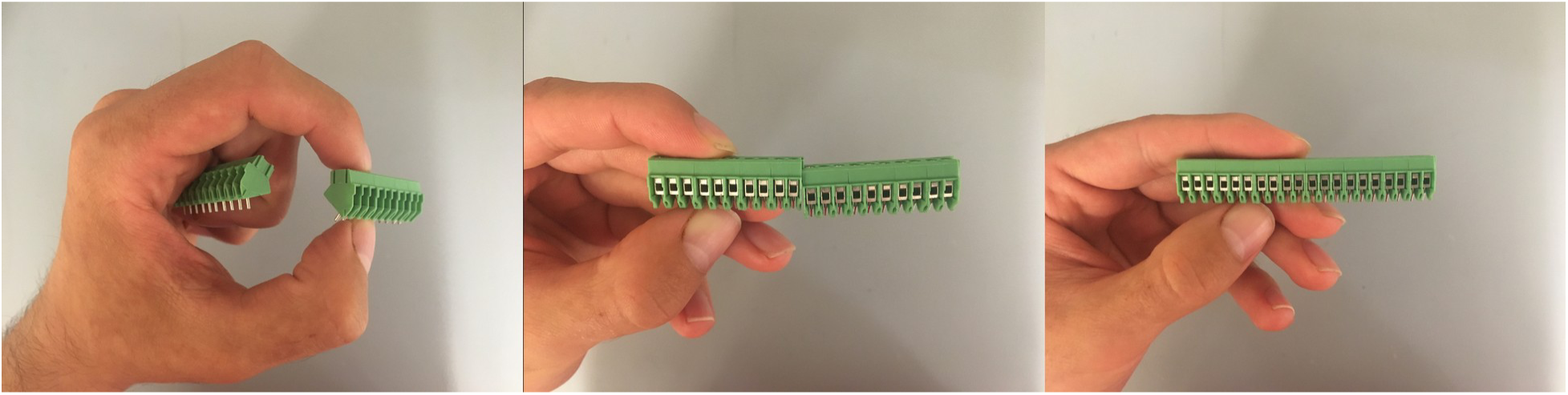
Demonstration of how to join terminals with interlocking sides.
6. Solder the terminal blocks and power jack on the top side of the board with a soldering iron and solder wire.
7. Solder the stackable Arduino Mega headers taking care to install them all perpendicular to the board: if they point at different angles from each other, the shield will be very difficult to plug into an Arduino.

- First, solder one pin of each header. Then align the shield to an Arduino Mega and adjust misaligned headers by reheating the single soldered joint. Make sure that the shield easily plugs into the Arduino before soldering the rest of the pins. See reference [23] for a full tutorial.
8. Solder the capacitor (take care to install with the correct polarity) and then clip the leads.
9. Plug the KATARA shield into an Arduino Mega (Figure 6).

**Figure 6.**
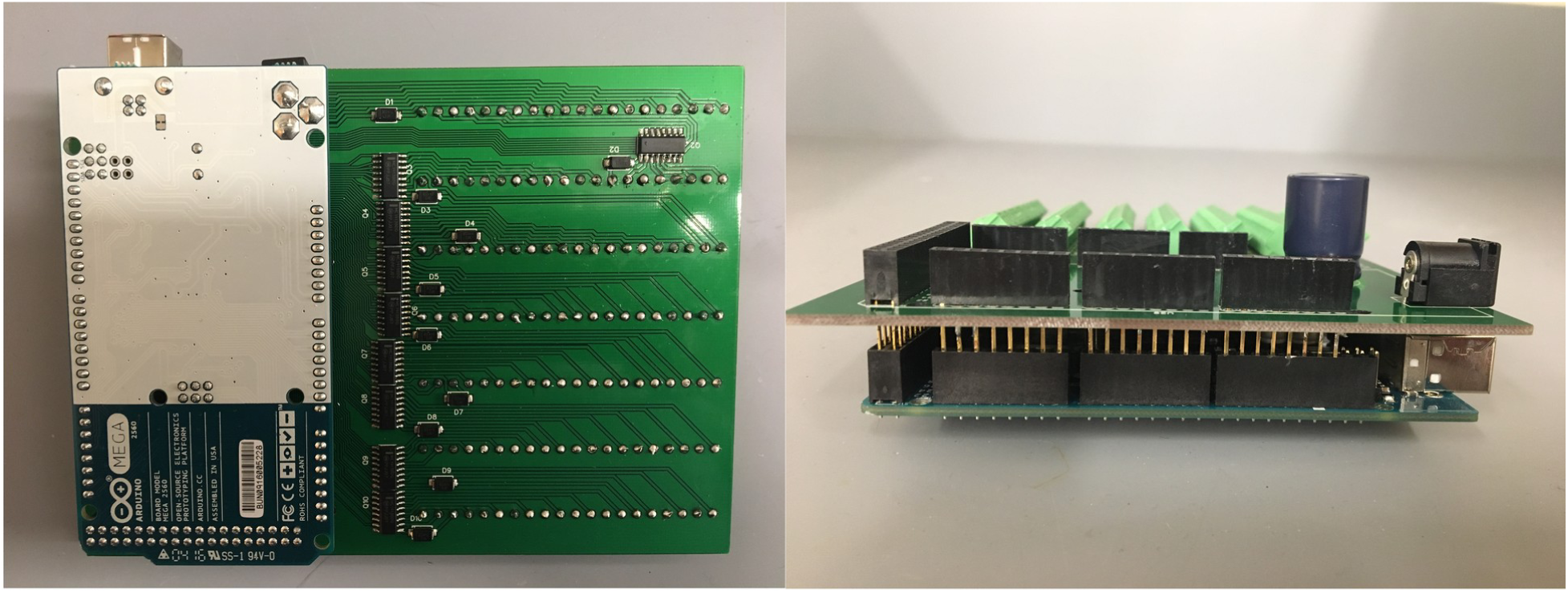
A KATARA Shield plugged into an Arduino Mega.
10. Attach solenoid valve lead wires to the terminal blocks. Clamp the leads in place by tightening the screws when the board is not powered to avoid shocking hazard (Figure 7).

**Figure 7.**
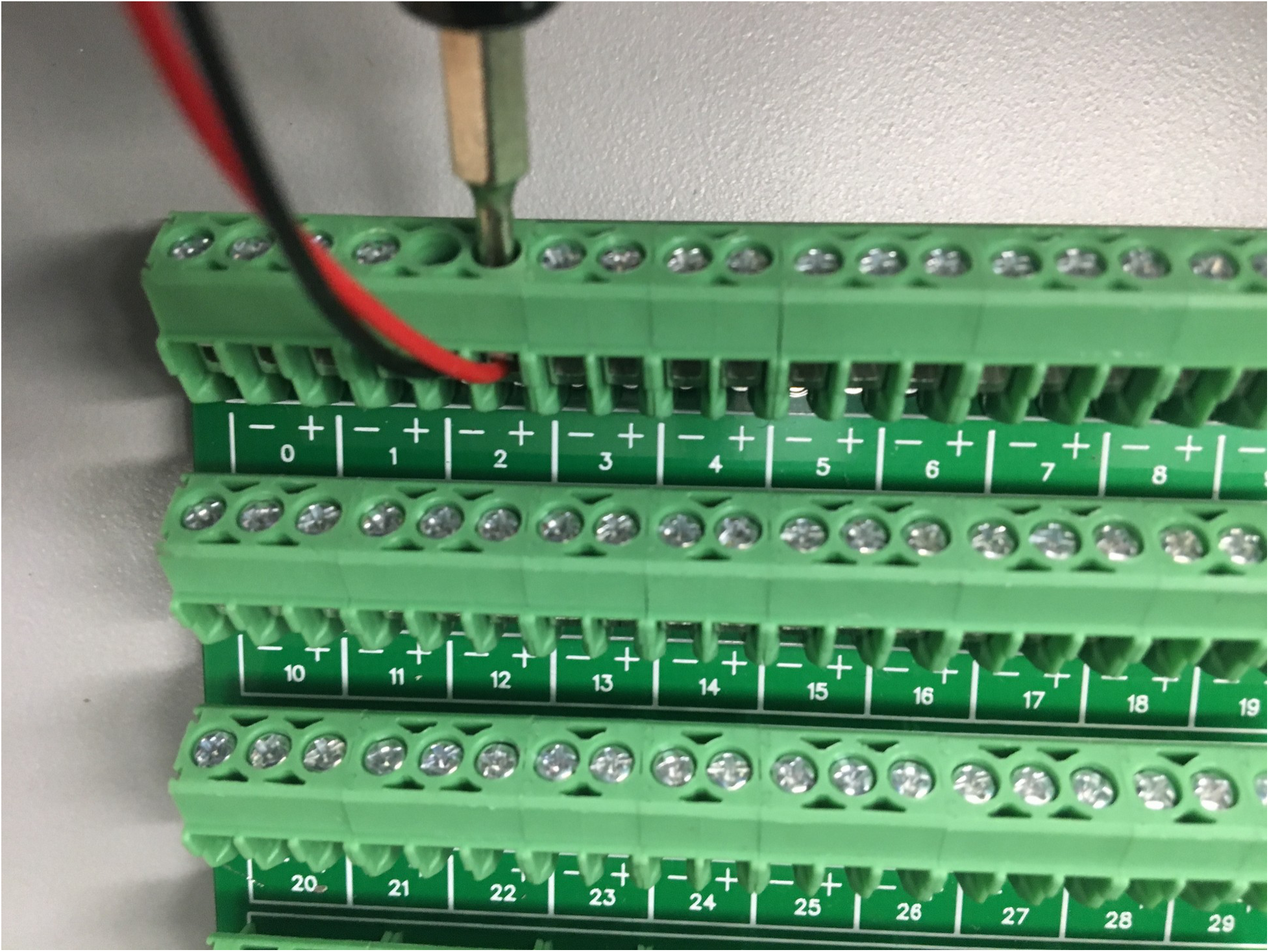
Solenoid valve lead wires can be clamped into place by tightening the screws on the terminal blocks.

## 6 Operation Instructions

To operate the KATARA Shield, first install the KATARA firmware to the Arduino using the Arduino IDE. The KATARA python software requires python 2.7 and the pyserial package. To run the GUI, open a terminal window and navigate into the KATARA_Software folder (available on Github), then run the command: python main.py

Once the software is open and connected to the Arduino, the user may open and close valves manually, specify pump modules, edit and save protocols, and load saved protocols as custom control buttons. For more detailed instructions on how to use and install the KATARA Firmware and Software, see the supplement.

## 7 Hardware validation

To evaluate the performance of the KATARA amplifier circuit, we measured the voltage between the collector of a Darlington amplifier and ground with an oscilloscope (TBS 1052B-EDU Tektronix) as the circuit switched off a solenoid valve (Figure 8). Figure 8 shows that before switching off at time zero, the digital output from the Arduino at the amplifier base is high and the voltage at the collector is zero. At time zero when the Arduino’s signal to the transistor base goes low, the amplifier circuit stops conducting. The voltage at the collector then spikes as the solenoid continues to drive current, but plateaus when the voltage across the Zener diode reaches its breakdown level. The high collector voltage reverses the current through the solenoid, then drops below the Zener breakdown level after about one millisecond and decays to the source voltage within another two milliseconds (Figure 8). This demonstrates that the electrical response of the KATARA circuit connected to Pneumadyne S10MM-31-24-2 valves is under three milliseconds, which is less than its specified ten millisecond de-energization time and on the same order as a microfluidic valve’s response time [1]. Finally, to demonstrate that the circuit is capable of controlling a microfluidic device with integrated pumps, we used the KATARA control system to drive a peristaltic pumping sequence (Video 1).

**Figure 8.**
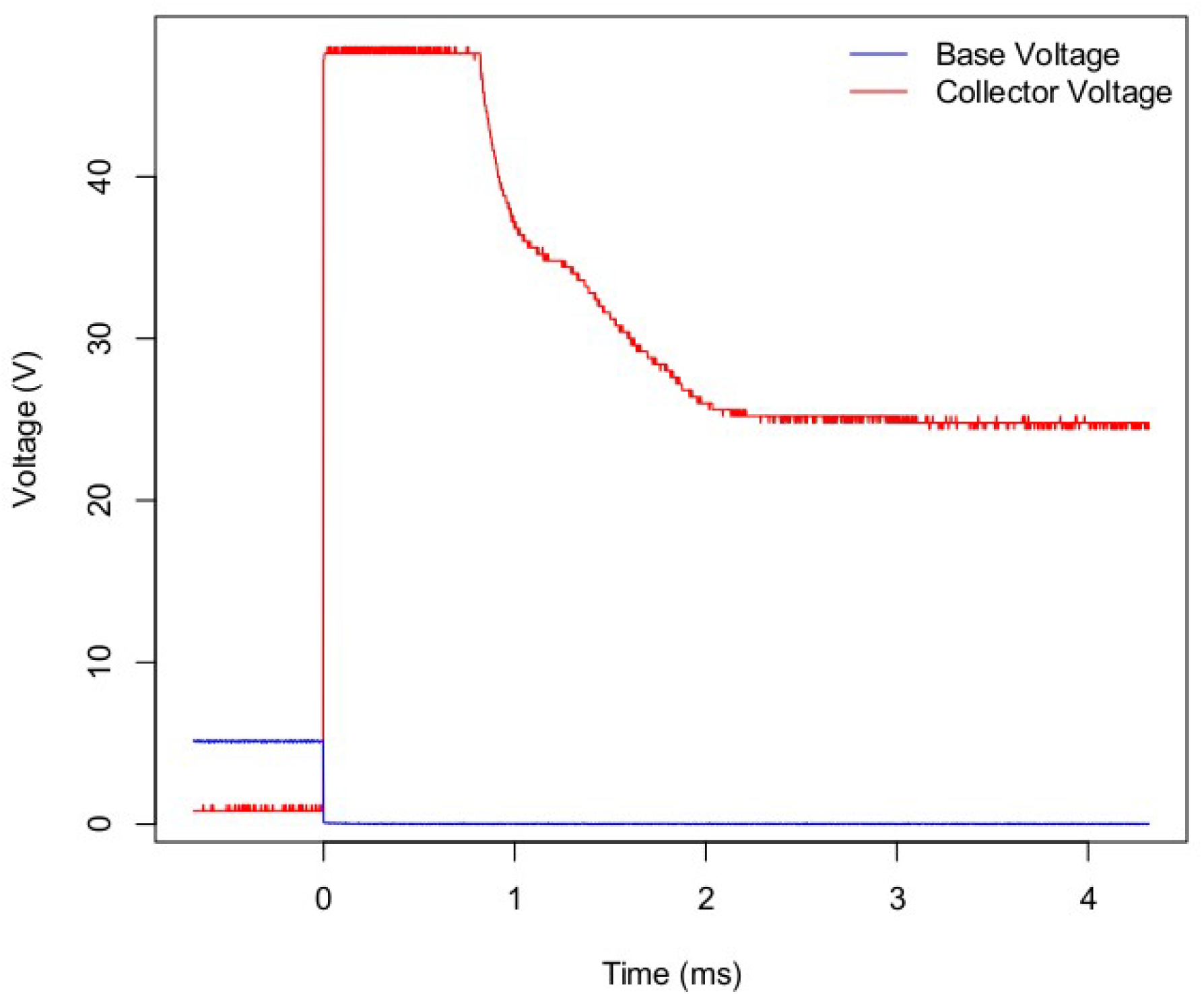
The electrical response of Pneumadyne S10-31-23-2 solenoid valves when the KATARA shield switches them off (at time zero) takes less than three milliseconds.

## Discussion

The KATARA provides a user-friendly solution to control solenoid valves at low cost. A complete microfluidic platform also includes pneumatic infrastructure to relay pressure to the solenoid valves and operate the microfluidic device. In this issue, Brower et al. present a comprehensive pneumatic platform for microfluidic large-scale integration[16]. The KATARA may be used as an alternate control module in this platform, as it serves as a low-cost alternative to the Wago controller. The KATARA shield also has the capability to control microfluidic devices remotely without a computer. This opens up the possibility to use microfluidic devices with integrated valves in field settings. The KATARA shield also maintains the Arduino’s ability to use its digital pins for purposes other than driving solenoid valves when valves are not connected to their amplifying circuits. Additionally, the KATARA may be suitable for purposes other than microfluidics including soft robotics, driving motors, and powering light sources. As we continue to develop the KATARA, we will post hardware and software updates on the Streets Lab website (http://streetslab.berkeley.edu/tools/katara/).

## Acknowledgements

We would like to thank Nicolas Altemose and Andre Lai for their help in preparing microfluidic devices to demonstrate the KATARA control system. We also would like to thank them, Anushka Gupta, and Gabriel Dorlhiac for their helpful comments on the manuscript. This work was supported by the University of California, Berkeley Department of Bioengineering. Aaron Streets is a Chan Zuckerberg Investigator.

## References

[1] M.A. Unger, H. P. Chou, T. Thorsen, A. Scherer, S. R. Quake, Monolithic Microfabricated Valves and Pumps by Multilayer Soft Lithography. Science, 228(5463), (2012) 113–116. 10.1126/science.288.5463.113

[2] T. Thorson, S. J. Maerkl, S. R. Quake, Microfluidic Large-Scale Integration. Science, 298(5593), (2002) 580–584. 10.1126/science.1076996

[3] J. Melin, S. R. Quake, Microfluidic Large-Scale Integration: The Evolution of Design Rules for Biological Automation. Annu Rev Biophys Biomol Struct, 36 (2007) 213–231. 10.1146/annurev.biophys.36.040306.132646

[4] C. L. Hansen, M. O. Sommer, S. R. Quake, Systematic investigation of protein phase behavior with a microfluidic formulator. Proc Natl Acad Sci USA, 101(40), (2004) 14431–14436. 10.1073/pnas.0405847101

[5] S. Kim, A. M. Streets, R. R. Lin, S. R. Quake, S. Weiss, D. S. Majumdar, High-throughput single-molecule optofluidic analysis. Nat Methods, 8(3), (2011) 242–245. 10.1038/nmeth.1569

[6] S. J. Maerkl, S. R. Quake, A systems approach to measuring the binding energy landscapes of transcription factors. Science, 315(5809), (2007) 233–237. 10.1126/science.1131007

[7] F. K. Balagaddé, L. You, C. L. Hansen, F. H. Arnold, S. R. Quake. Long-term monitoring of bacteria undergoing programmed population control in a microchemostat. Science, 309(5731), (2005) 137–140. 10.1126/science.1109173

[8] R. Gómez-Sjöberg, A. A. Leyrat, D. M. Pirone, C. S. Chen, S. R. Quake, Versatile, fully automated, microfluidic cell culture system. Anal Chem, 79(22), (2007) 8557–8563. 10.1021/ac071311w

[9] E. A. Otteson, J. W. Hong, S. R. Quake, J. R. Leadbetter, Microfluidic digital PCR enables multigene analysis of individual environmental bacteria. Science, 314(5804), (2006) 1464–1467. 10.1126/science.1131370

[10] J. L. Garcia-Cordero, S. J. Maerkl, A 1024-sample serum analyzer chip for cancer diagnostics, Lab on a Chip, 14(15) (2014) 2642–2650. doi:10.1039/c3lc51153g

[11] J. L. Garcia-Cordero, C. Nembrini, A. Stano, J. A. Hubbell, S. J. Maerkl, A high-throughput nanoimmunoassay chip applied to large-scale vaccine adjuvant screening, Integrative Biology, 5(4) (2013) 650–658. doi:10.1039/c3ib20263a

[12] Y. Marcy, C. Ouverney, E. M. Bik, T. Lösekann, N. Ivanova, H. G. Martin, E. Szeto, D. Platt, P. Hugenholtz, D. A. Relman, S. R. Quake, Dissecting biological “dark matter” with single-cell genetic analysis of rare and uncultivated TM7 microbes from the human mouth. Proc Natl Acad Sci U S A, 104(29), (2007) 11889–11894. 10.1073/pnas.0704662104

[13] A. M. Streets, X. Zhang, C. Cao, Y. Pang, X. Wu, L. Xiong, L. Yang, Y. Fu, L. Zhao, F. Tang, Y. Huang, Microfluidic single-cell whole-transcriptome sequencing. Proc Natl Acad Sci U S A, 111(19), (2014) 7048–7053. 10.1073/pnas.1402030111

[14] K. Brower, A. K. White, P. M. Fordyce, Multi-step Variable Height Photolithography for Valved Multilayer Microfluidic Devices. J Vis Exp, (110), (2017) 10.3791/55276

[15] R. Gómez-Sjöberg, USB-Based Controller. https://sites.google.com/site/rafaelsmicrofluidicspage/valve-controllers/usb-based-controller, (accessed 07.20.17)

[16] K. Brower, R. Puccinelli, C. Markin, T. Shimko, R. Garcia-Gomez, P. M. Fordyce, An Open-Source, Programmable Pneumatic Setup for Operation and Automated Control of Single‐ and Multi-layer Microfluidic Devices. BioRxiv, (2017) https://doi.org/10.1101/173468

[17] H. H. Strey, Open Hardware Microfluidics Controller Arduino Shield. https://streylab.com/blog/2015/4/8/open-hardware-microfluidics-controller-arduino-shield, 2015 (accessed 07.20.17)

[18] B. Li, L. Li, A. Guan, Q. Dong, K. Ruan, R. Hu, Z. Li, A smartphone controlled handheld microfluidic liquid handling system. Lab Chip, (20), (2014) 4085–4092. 10.1039/C4LC00227J

[19] F. Piraino, F. Volpetti, C. Watson, S. J. Maerkl, A Digital–Analog Microfluidic Platform for Patient-Centric Multiplexed Biomarker Diagnostics of Ultralow Volume Samples. ACS Nano, 10(1), (2016) 1699–1710. doi:10.1021/acsnano.5b07939

[20] N. Seidle, Solder Paste Stenciling. https://www.sparkfun.com/tutorials/58, 2006 (accessed 07.20.17)

[21] ON Semiconductor, Soldering and Mounting Techniques: Reference Manual. https://www.onsemi.com/pub/Collateral/SOLDERRM-D.PDF, 2016 (accessed 07.20.17)

[22] N. Seidle, Reflow Skillet. https://www.sparkfun.com/tutorials/59#Skillet, 2006 (accessed 07.20.17)

[23] Jimb0, Arduino Shields: Installing Headers (Assembly). https://learn.sparkfun.com/tutorials/arduino-shields/installing-headers-assembly, (accessed 07.20.17)

